# Human cerebrospinal fluid single exosomes in Parkinson’s and Alzheimer’s diseases

**DOI:** 10.1101/2023.12.22.573124

**Authors:** Koya Yakabi, Eloise Berson, Kathleen S. Montine, Sean C. Bendall, Michael J. MacCoss, Kathleen L. Poston, Thomas J. Montine

**Affiliations:** Department of Pathology, Stanford University, Stanford, CA, USA; Department of Genome Sciences, University of Washington, Seattle, WA, USA; Department of Department of Neurology & Neurological Sciences, Stanford University, Stanford, CA, USA

## Abstract

Exosomes are proposed to be important in the pathogenesis of prevalent neurodegenerative diseases. We report the first application of solid-state technology to perform multiplex analysis of single exosomes in human cerebrospinal fluid (CSF) obtained from the lumbar sac of people diagnosed with Alzheimer’s disease dementia (ADD, n=30) or Parkinson’s disease dementia (PDD, n=30), as well as age-matched health controls (HCN, n=30). Single events were captured with mouse monoclonal antibodies to one of three different tetraspanins (CD9, CD63, or CD81) or with mouse (M) IgG control, and then probed with fluorescently labeled antibodies to prion protein (PrP) or CD47 to mark neuronal or presynaptic origin, as well as ADD- and PDD-related proteins: amyloid beta (Aβ), tau, α-synuclein, and Apolipoprotein (Apo) E. Data were collected only from captured events that were within the size range of 50 to 200 nm. Exosomes were present at approximately 100 billion per mL human CSF and were similarly abundant for CD9+ and CD81+ events, but CD63+ were only 22% to 25% of CD9+ (P<0.0001) or CD81+ (P<0.0001) events. Approximately 24% of CSF exosomes were PrP+, while only 2% were CD47+. The vast majority of exosomes were surface ApoE+, and the number of PrP-ApoE+ (P<0.001) and PrP+ApoE+ (P<0.01) exosomes were significantly reduced in ADD vs. HCN for CD9+ events only. Aβ, tau, and α-synuclein were not detected on the exosome surface or in permeabilized cargo. These data provide new insights into single exosome molecular features and highlight reduction in the CSF concentration of ApoE+ exosomes in patients with ADD.

## INTRODUCTION

Exosomes are a subset of extracellular vesicles (EVs) produced by cells and that are characterized by plasma membrane-bound molecules that direct their trafficking and cytoplasmic cargo. Multiple groups have investigated exosomes from a variety of body fluids in relationship to neurodegenerative diseases, and these have used a variety of bulk preparation techniques that enrich exosomes from other EVs and lipoproteins with each method having comparative advantages and disadvantages [1,2]. Some groups have attempted to use flow cytometric techniques to probe single exosomes or EVs, but it remains unclear to us and others whether particles of this small size can be isolated reproducibly by current flow technologies [1]. Here, we report the first application of solid-state technology to perform multiplex analysis of single exosomes in human cerebrospinal fluid (CSF) from people diagnosed with Parkinson’s disease with dementia (PDD) or Alzheimer’s disease dementia (ADD), or who were age-matched healthy controls (HCN).

## METHODS

CSF samples were obtained from 90 individuals enrolled in research cohorts at Stanford University. All participants (or representatives) provided written informed consent; all protocols and assessments were performed with approval by the institutional internal review board. Diagnoses of ADD (n=30) or PDD (n=30) were made using current consensus criteria [3,4]. Healthy controls (HCN, n=30) also were enrolled and evaluated in the same research cohorts. Lumbar CSF samples from these 90 individuals (**Table 1**) were thawed once, aliquoted, refrozen at -80°C, and then thawed only once more for analysis. Additional aliquots from each group were combined to create a pooled sample that was used as a technical control for each run.

**Table 1.** Characteristics of research participants.

|  | All Samples<br>*n*=90 | ADD *n*=30 | PDD *n*=30 | HNC *n*=30 |
| --- | --- | --- | --- | --- |
| Age (Mean-SD) | 70.0 SD=7.7 | 69.9 SD=7.6 | 71.4 SD=7.0 | 68.8 SD=8.5 |
| Gender # of male (% male) | *n*=51 (56.7%) | *n*=16 (53.3%) | *n*=19 (63.3%) | *n*=16 (53.3%) |
| ApoE ε4 (% ε4 allele) | 39/146 (26.7%) | 26/48 (54.2%) | 8/56 (14.3%) | 5/42 (11.9%) |

CSF samples were analyzed using Leprechaun from Unchained Labs (Pleasanton, CA), which utilizes a solid-state technology to immunocapture EVs with antibodies to tetraspanins, measure size of a single particle by interferometry, and phenotype each particle using immunofluorescence microscopy. CSF was used neat for the data reported; however, all antibodies for immunofluorescence microscopy also were tested at 1:2 and 1:10 dilutions of CSF to ensure the expected reduction in signal. Each CSF sample was analyzed on four different capture panels designed to capture single events with mouse monoclonal antibodies to one of three different tetraspanins that are characteristic of exosomes (CD9, CD63, or CD81) or with mouse (M) IgG control in duplicate or triplicate. Data were collected only from captured events that were within the size range of 50 to 200 nm.

Particles within the appropriate size range on all four capture panels were probed with fluorescently tagged antibodies, conjugated using Thermo Fisher Alexa Fluor Kits (AF-488 catalog #A20181, AF-555 catalog #A20187, AF-647 catalog #A20186) to PrP (3F4) (BioLegend catalog #800301) or CD47 (MIAP410) (InVivoMAb catalog #BE0283), which are highly enriched in neuronal membranes and presynaptic membranes, to assess neuronal origin of the captured particle. These data were averaged across the four capture panels such that a single average value was calculated for each individual CSF sample. One capture panel each also was probed with tagged antibodies to disease-associated proteins: BioLegend Amyloid Beta (6E10) (catalog # 803001), Millipore Sigma Tau (Tau-5) (catalog # MABN162), BioLegend Alphα-Synuclein (LB509) (catalog #807701), or BioLegend ApoE (D6E10) (catalog #803301) . We ran 16 chips per session: 5 with HNC samples, 5 with ADD samples, 5 with PDD samples, and one with the pooled technical control. Pooled technical control was applied in the same position on each chip. Samples from the diagnostic groups were randomly distributed across the chip array.

Isolated exosomes were also subjected to cargo analysis. The analysis involved following the manufacturer’s cargo protocol for internal cargo assessment and, in parallel, applying the surface protocol as a control.

### Antibodies

PrP (3F4) (BioLegend catalog #800301), CD47 (MIAP410) (InVivoMAb catalog #BE0283), CD56 (NCAM16.2) (Fischer Scientific catalog #BDB559043), GLAST/EAAT (polyclonal) (Invitrogen catalog #PA5-72895), Aquaporin-4 (E5206) (BioLegend catalog #600352), CD11b (M1/70) (BioLegend catalog #101247), CD68 (D4B9C) (Cell Signaling Technology catalog #76437S), Amyloid Beta (6E10) (BioLegend catalog # 803001), Tau (Tau-5) (Millipore Sigma catalog # MABN162), Alphα-Synuclein (LB509) (BioLegend catalog #807701), or ApoE (D6E10) (BioLegend catalog #803301).

## RESULTS

Characteristics of individuals in the four groups are summarized in **Table 1**. We used lumbar CSF samples from individuals in these three diagnostic groups to capture single exosomes using three different tetraspanin antibodies (mouse immunoglobulin (MIgG) as negative control) and limited their size to between 50 and 200 nm (**Figure 1**).

**Figure 1.**
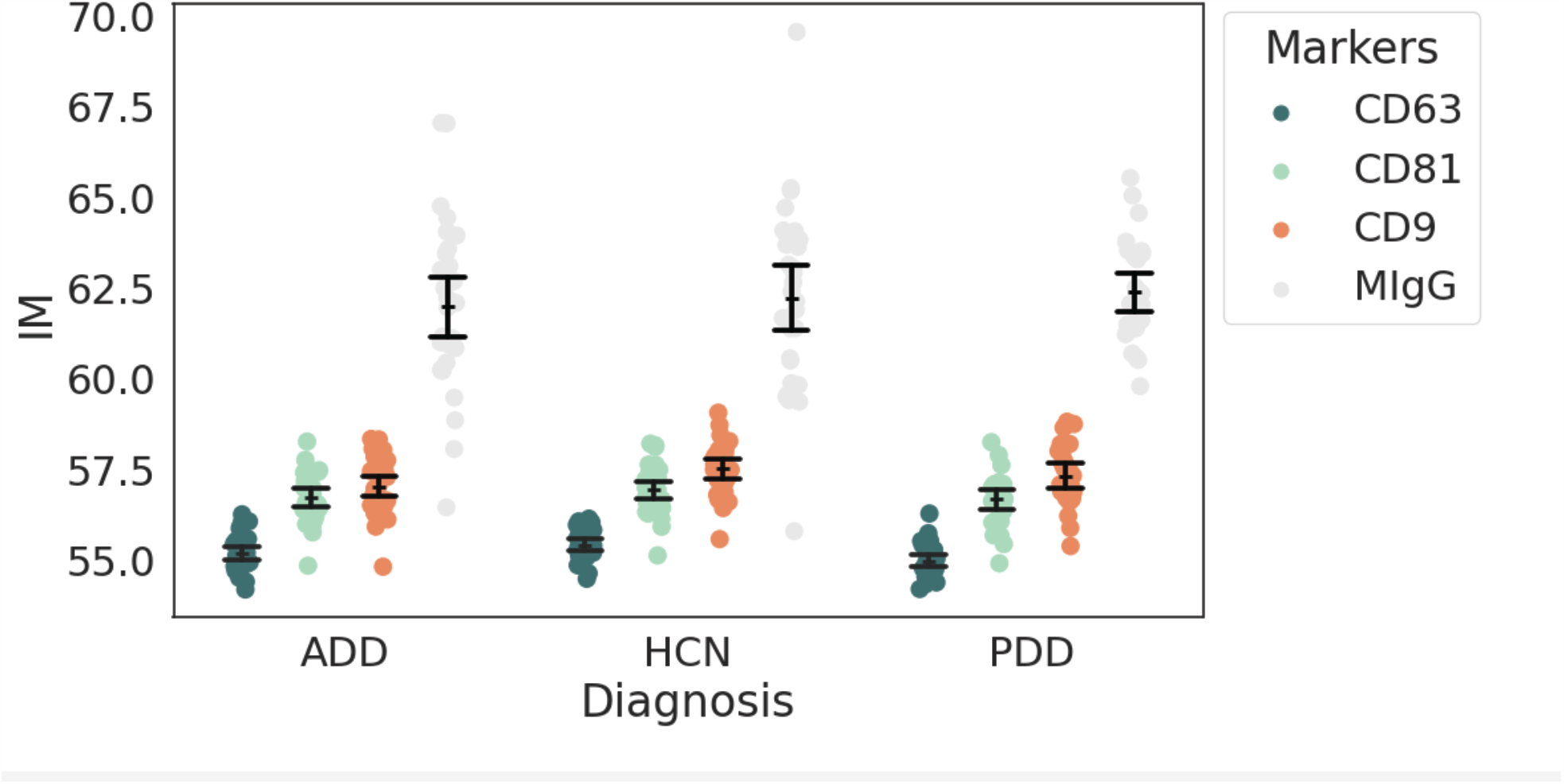
Single event size distribution measured by interferometry (microns) for each tetraspanin capture antibodies and mouse immunoglobulin (MIgG) negative control stratified by diagnosis groups: Alzheimer’s disease dementia (ADD), Parkinson’s disease dementia (PDD), healthy controls (HCN).

There were 1.25 ± 0.29, 1.35 ± 0.31, and 1.33 ± 0.28 x10^8^ total exosomes per mL human CSF in the ADD, HCN, and PDD groups, with no significant difference among diagnostic groups. Two-way ANOVA for number of exosomes stratified by capture antibody and diagnostic group was statistically significant only for capture antibody (P<0.0001) but not diagnostic group or interaction between the two factors. Tukey’s corrected multiple pairwise comparisons showed that CD63+ exosomes were 22% to 25% of CD9+ (P<0.0001) or CD81+ (P<0.0001) exosomes in each diagnostic group; abundance of CD9+ or CD81+ exosomes were not different in any of the diagnostic groups (**Figure 2**).

**Figure 2.**
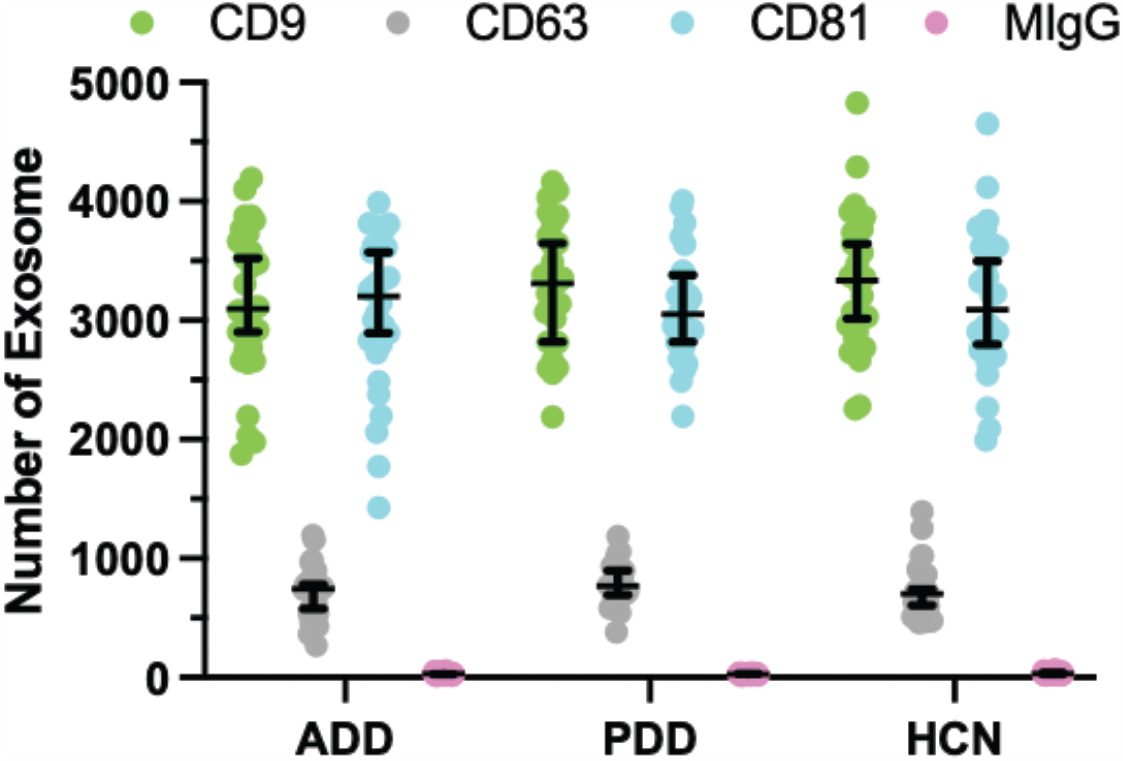
Number of exosomes in 20 μL of human CSF from individuals diagnosed with Alzheimer’s disease dementia (ADD), Parkinson’s disease dementia (PDD), or healthy controls (HCN). Aliquots from each CSF sample were separately captured by mouse monoclonal IgG to CD9, CD63, or CD81 or to mouse IgG (MIgG); captured single events were restricted in size to 50 to 200 microns. Number of exosomes stratified by capture antibody and diagnostic group was statistically significant only for tetraspanin capture antibody (P<0.0001). CD63+ exosomes were significantly less abundant than CD9+ (P<0.0001) or CD81+(P<0.0001) exosomes in each diagnostic group.

We tried multiple antibodies to neuron-specific surface antigens to detect neuronal exosomes, including PrP, CD47, and NCAM. Of these, an antibody to PrP yielded the most reproducible estimate of the largest fraction of exosomes. Using this antibody, approximately 24% of all CSF exosomes were PrP+, and presumably enriched for neuron-derived exosomes. When stratified by tetraspanin capture antibody and diagnostic group, we again observed significantly fewer CD63+PrP+ events (P<0.0001 for both) than either CD9+PrP+ or CD81+PrP+ events, and CD9+PrP+ and CD81+PrP+ events were not different (**Figure 3A**). The same analysis for PrP-events revealed the same relationships with CD63+PrP-events being less abundant (P<0.0001 for both) than either CD9+PrP- or CD81+PrP-events, and CD9+PrP- and CD81+PrP-events not different (**Figure 3B**).

**Figure 3.**
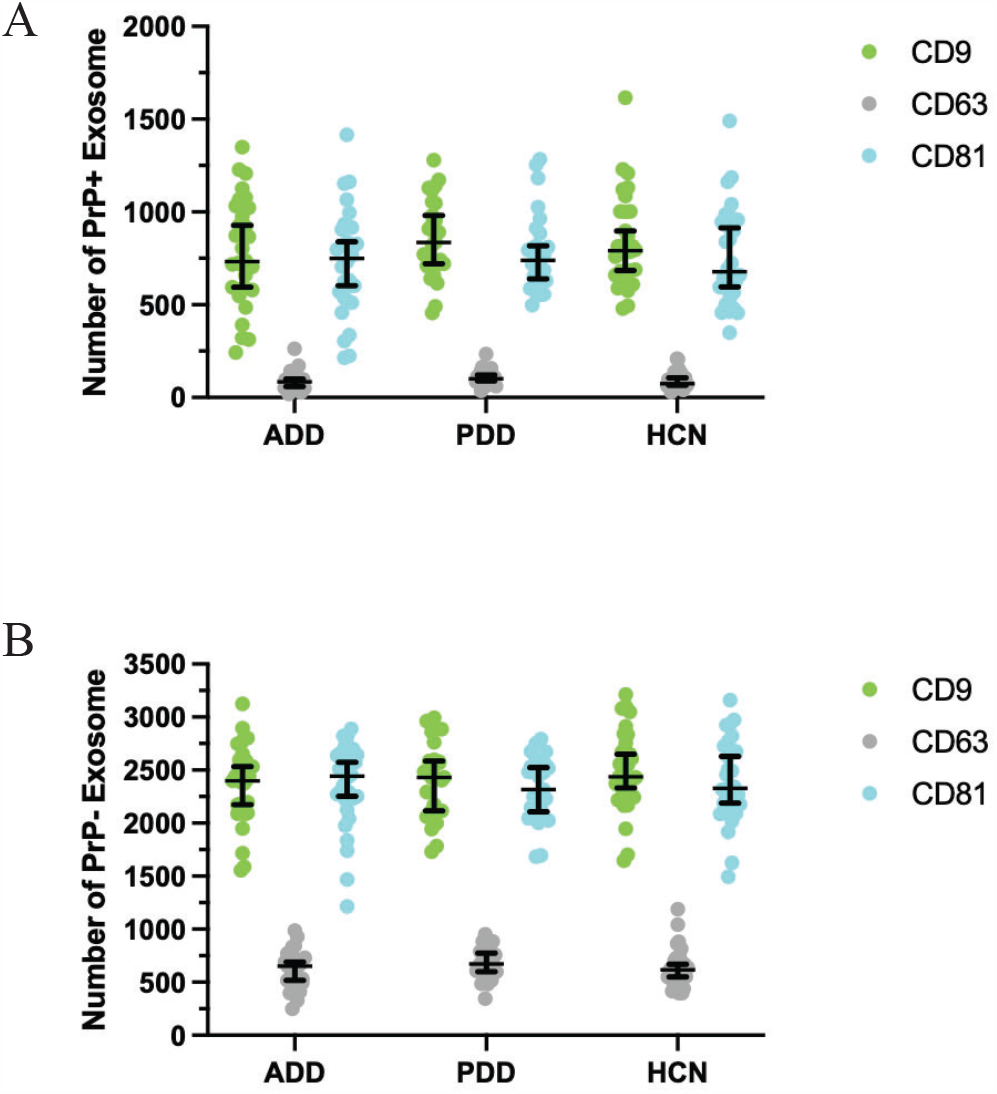
Number of PrP+ (**A**) and PrP-(**B**) exosomes in 20 μL of human CSF from individuals diagnosed with Alzheimer’s disease dementia (ADD), Parkinson’s disease dementia (PDD), or healthy controls (HCN). Aliquots from each sample were separately captured by mouse monoclonal IgG to CD9, CD63, or CD81 or to mouse IgG (MIgG) and captured single events restricted in size to 50 to 200 microns.

We further subcategorized PrP+ exosomes with CD47, a highly expressed plasma membrane bound pre-synaptic protein. Less than 2% of all the CSF exosomes were CD47+; however, the relationship of CD47+ exosomes to diagnostic groups was not different from PrP+ exosomes (not shown). We also tried multiple antibodies to astrocytic (GLAST, Aquaporin-4) or microglial (CD11b, CD68) surface antigens, but none yielded detectable signal in human CSF.

### Disease-related proteins

While tetraspanin capture, PrP, and CD47 were used on all panels, we varied one antibody on each panel to evaluate proteins related to PDD or ADD: Aβ, tau, α-synuclein, or ApoE. We observed no signal for Aβ, tau, or α -synuclein. In contrast, we observed abundant ApoE+ exosomes (**Figure 4**). Indeed, ApoE reactivity was present on the vast majority of exosomes, and followed the same pattern for tetraspanin capture as observed above for total exosomes with CD9+ApoE+ (**Figure 4A**), similar to CD81+ApoE+ (**Figure 4B**), and both more concentrated than CD63+ApoE+ exosomes (**Figure 4C**); there was no significant difference among diagnostic groups.

**Figure 4.**
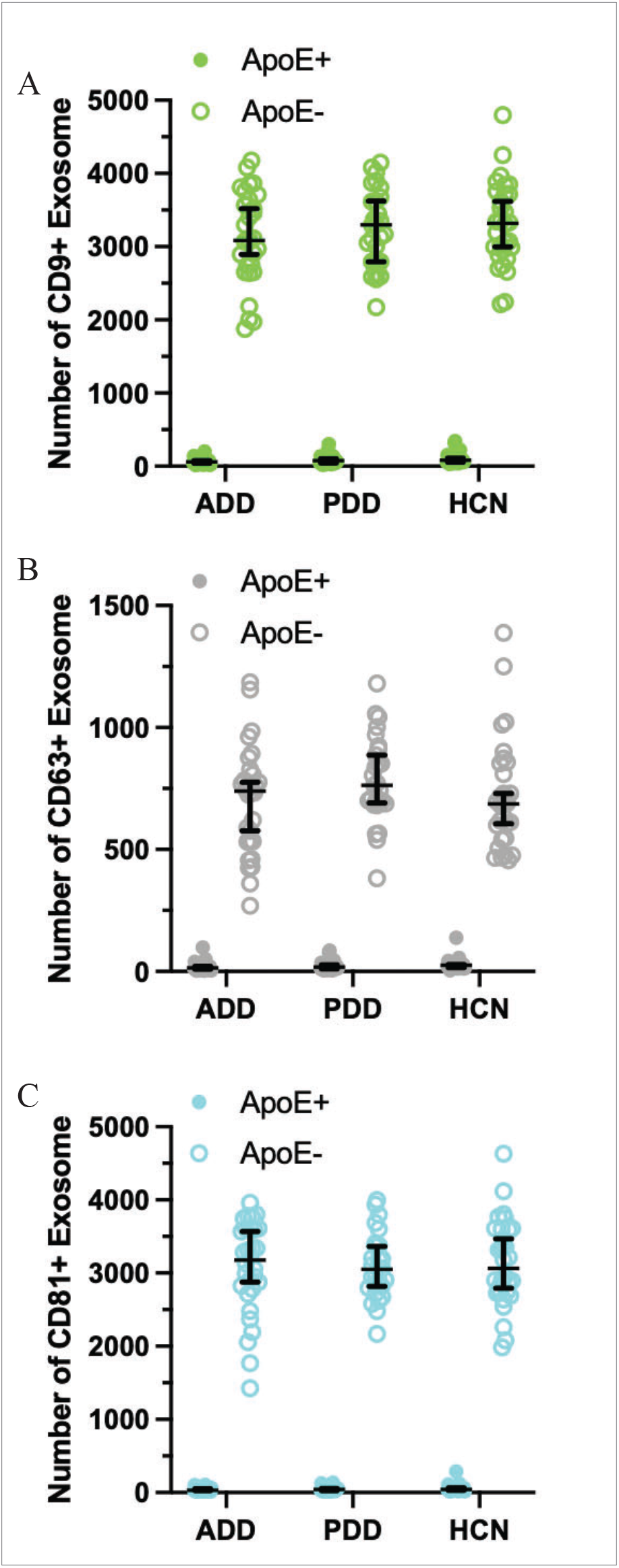
Number of ApoE+ and ApoE - exosomes in 20 μL of CSF from individuals diagnosed with AD dementia (ADD), Parkinson’s disease dementia (PDD), or healthy controls (HCN). Aliquots from each CSF sample were separately captured by mouse monoclonal IgG to CD9 (**A**), CD63 (**B**), or CD81(**C**) and restricted in size to 50 to 200 microns.

We next analyzed the expression of ApoE on exosomes derived from neurons (PrP+) or not (PrP-) **(Figure 5**). The number of PrP-ApoE+ (**Figure 5A**, P<0.001) and PrP+ApoE+ (**Figure 5B**, P<0.01) exosomes were significantly reduced in ADD vs. HCN for CD9+ events only.

**Figure 5.**
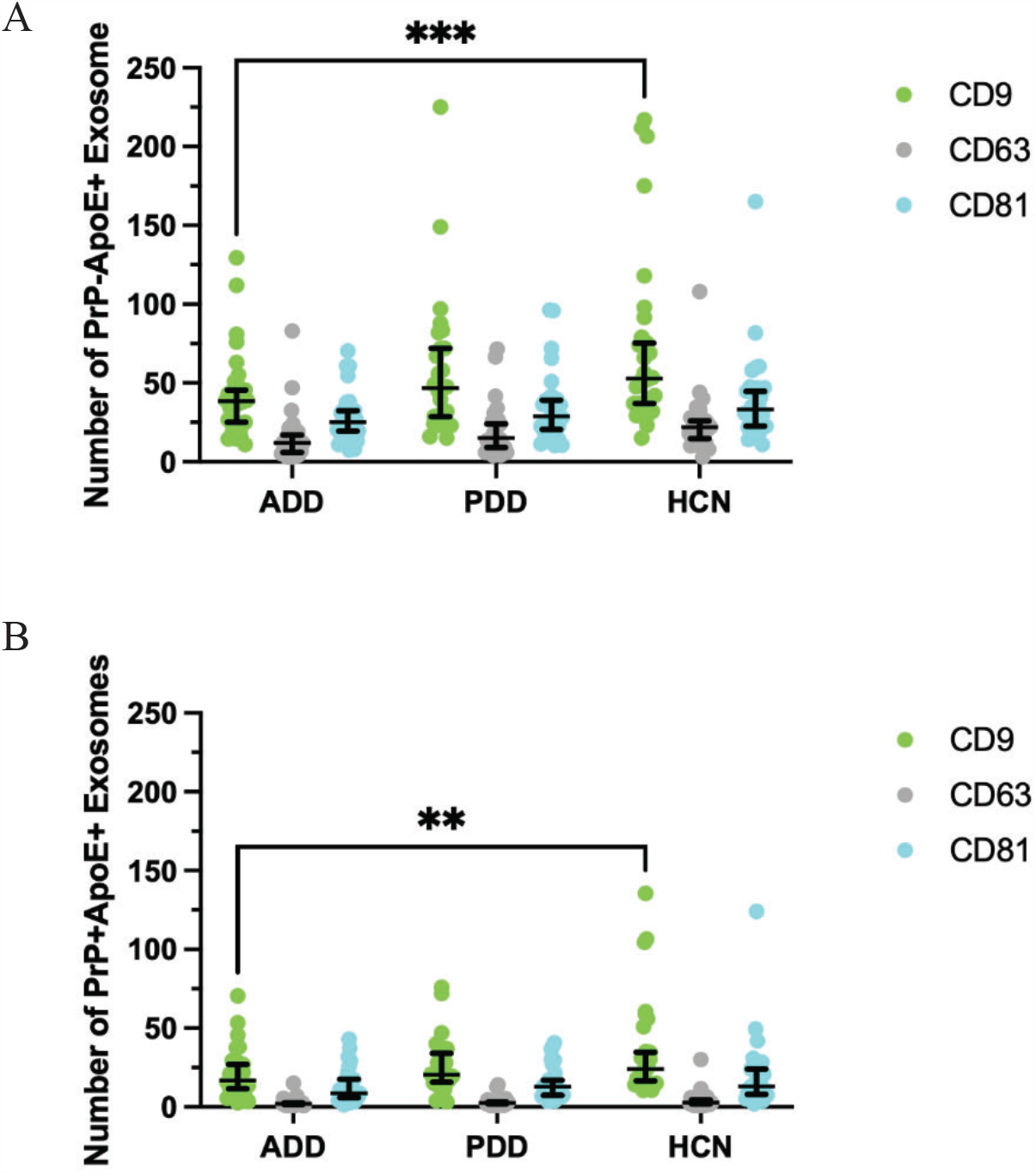
Number of PrP-ApoE+ (**A**) and PrP+ApoE+ (**B**) exosomes in 20 μL of human CSF from individuals diagnosed with Alzheimer’s disease dementia (ADD), Parkinson’s disease dementia (PDD), or healthy controls (HCN). Aliquots from each CSF sample were separately captured by mouse monoclonal IgG to CD9, CD63, or CD81 and restricted in size to 50 to 200 microns. The number of PrP-ApoE+ (P<0.001) and PrP+ApoE+ (P<0.01) exosomes was significantly reduced in ADD vs. HCN for CD9+ events only.

The tetraspanins, PrP, CD47, α-synuclein, and ApoE all bind to the plasma membrane and, as expected, were not detected in greater amounts following permeabilization to detect both exosome surface and cargo proteins. Contrary to expectation, we also did not detect additional Aβ or tau protein following permeabilization, indicating that these two proteins are neither detectable on the surface nor in cargo of CSF exosomes assayed by this method.

## DISCUSSION

Exosomes, a subclass of EVs, are elaborated by cells, bound by a fragment of the plasma membrane, and harbor cytoplasmic cargo. Here we tested the hypothesis that the concentration and molecular composition of exosomes was altered by PDD or ADD. We used neat CSF to avoid the now well-known confounding effects of different methods for enriching exosomes [5,6]. We used a solid-state method for detecting individual EVs with the characteristic molecular features and size of exosomes. Overall, we estimated that the concentration of exosomes in human CSF is approximately 100 million per mL.

Our experiments focused on proteins detectable on the surface of exosomes and not on their cargo. Following several pilot experiments, we settled on PrP, a pan neuronal protein bound to the external face of the plasma membrane, as a marker for neurons and CD47, a highly abundant presynaptic membrane spanning protein with defined extracellular domain, as a marker for presynaptic derivation. In addition to the tetraspanins, we used antibodies to these two proteins in all four panels. We also used an antibody to four different disease-associated proteins, a different antibody on each of the four panels: Aβ, tau, α-synuclein, and ApoE. α-synuclein is a presynaptic protein that is part of the synaptic vesicle apparatus and tau is a microtubule-associated protein; neither has a known extracellular domain. Aβ is an endoproteolytic product of APP, which is a single plasma membrane-spanning protein with an extracellular domain. Some but not all data support that Aβ peptides bind to the extracellular face of plasma membranes. We included antibodies to these three disease-associated proteins because each has been used as CSF biomarker for AD or PD with the presumption that they are floating free in CSF. We tested the hypothesis that Aβ, tau, or α-synuclein may be bound to the surface of human CSF exosomes and found no detectable signal. It is entirely possible that any or all of these disease-associated proteins are part of exosome cargo; however, their absence from the surface of exosomes challenges the proposal that any of them are directing exosome trafficking.

ApoE in brain is predominantly translated in glia but also some neurons; it is a secreted protein that can bind to the extracellular face of the plasma membrane through a secretion-capture mechanism. We detected ApoE on the surface of CSF exosomes, predominantly in PrP-exosomes, aligning with its translation primarily in glia. Non-neuronal exosomes with ApoE (PrP-ApoE+) were significantly reduced in ADD vs. HNC for CD9 capture antibody but were not different between PDD and HCN. These data validate studies of human CSF that have reported reduced CSF ApoE concentration in patients with ADD and extend these finding to CSF CD9+ exosomes. Neuron-derived exosomes with surface ApoE (PrP+ApoE+) were less common and only significantly reduced in ADD vs HCN for CD9+ exosomes, again with no difference for PDD vs HCN.

There are limitations to our study. While we deliberately compared two different forms of dementia (ADD and PDD) to HCN for potentially confounding non-specific features of dementia, like decreased activity and altered behavior, these two diseases usually are treated differently. For this reason, we avoided directly comparing ADD to PDD but recognize that treatments may have impacted the comparisons to HCN. Some investigators have proposed that the lower CSF ApoE concentration in people with ADD is due to an effect of the *APOE* ε4 allele on transcription. Although well powered for biomarker determination, our sample size is underpowered to determine associations with *APOE* genotype. Therefore, our inability to detect an *APOE* genotype effect should be interpreted as null rather than negative. Finally, our data suggest CD9-specific impact on CSF exosome biology in people with ADD. We are unaware of any data that may inform this observation in human samples and await other experimental studies to shed light on this intriguing finding in human CSF samples from people with ADD.

## ACKNOWLEDGMENTS

This work was funded by AG065156, AG066515, and AG077443. The authors declare no conflicts of interest.

